# 5-Azacytidine alleviates Liver Fibrosis via modulation of TET1 and TET3 levels and restoration of global 5hmC

**DOI:** 10.64898/2026.11.18.767774

**Authors:** Venkatapathi Challa, Rachiraju H. Sindhuja, Truptimayee Barik, Arushi Mansingh, Sivapriya Pavuluri, Thomas M. Shiju, Kalyani Pathuri, Chaturvedula Tripura

## Abstract

DNA methylation and demethylation processes play a pivotal role in the onset and progression of liver diseases. The ten-eleven translocase enzymes (TET1, TET2 and TET3) are key DNA demethylases that oxidise 5-methylcytosine (5mC) to 5-hydroxymethylcytosine (5hmC), thereby regulating the cellular epigenetic state. Dysregulated TET expression has been increasingly associated with hepatic pathology. In this study, we assessed the DNA hydroxymethylation status of fibrotic and regenerating livers and correlated these changes with TET protein expression in a CCl₄ induced murine model of liver fibrosis. Fibrosis progression was accompanied by a progressive, stage-wise reduction in hydroxymethylation, assessed by 5hmC immunostaining, together with a corresponding decrease in TET1 and increase in TET3 protein levels detected by immunoblotting. TET1 immunostaining correlated with the pattern of 5hmC immunostaining in fibrotic liver tissue. Treatment with the epigenetic agent 5-Azacytidine (5-AZA) improved liver morphology and lowered serum AST and ALT activities. 5-AZA treatment restored 5hmC levels and TET1 expression while downregulating TET3 in total liver tissue. Complementary *in vitro* studies in mouse primary and human stellate cells (LX-2) corroborated an increase in TET3 that correlated with elevated 5hmC, both of which were attenuated in stellate cells treated with 5-AZA. The reduction in TET3 corresponded with decreased expression of fibrogenic markers; α-smooth muscle actin and Collagen-I. Collectively, this study identifies a progressive loss of 5hmC during liver fibrosis and demonstrates that pharmacological modulation via an epigenetic agent restores 5hmC homeostasis and TET-associated hydroxymethylation, attenuating fibrotic remodelling. These findings support targeting the DNA methylation-demethylation axis, and TET regulation in particular, as a promising therapeutic avenue for liver fibrosis.

## 1. Introduction

Chronic liver disease (CLD) is characterized by progressive events of hepatic injury, inflammation, and replacement of normal parenchyma with fibrotic scar tissue, ultimately leading to liver failure [1]. Liver injury triggers activation of the quiescent vitamin A storing hepatic stellate cells (HSCs), which transdifferentiate into proliferative myofibroblast-like cells that secrete excessive extracellular matrix (ECM) components, predominantly collagen. This process drives liver fibrosis, for which effective pharmacological interventions remain limited [2,3]. Mounting evidence implicates epigenetic remodelling as a major driver of HSC activation and fibrogenesis. Activated HSCs display widespread alterations in DNA methylation and histone modifications, encompassing both hypo- and hypermethylation at promoter and CpG-rich regions [4–6]. Understanding how these epigenetic regulators are perturbed, and how they might be modulated to restore homeostasis, offers new opportunities for antifibrotic therapy. DNA methylation, marked by the addition of a methyl group to cytosine to form 5-methylcytosine (5mC), serves as a stable epigenetic modification that regulates gene silencing and chromatin structure. Conversely, 5-hydroxymethylcytosine (5hmC) which is an oxidative and stable intermediate derivative of 5mC, serves as an indicator of active DNA demethylation [7,8]. The ten-eleven translocation (TET) family of dioxygenases TET1, TET2, and TET3, catalyze the stepwise sequential oxidation of 5mC to 5hmC, 5-formylcytosine (5fC), and 5-carboxylcytosine (5caC) [9,10]. The final removal of these oxidized cytosine derivatives and their replacement with unmethylated cytosine is mediated by thymine–DNA glycosylase (TDG) and the base excision repair (BER) pathway. This dynamic cycle is completed when DNA methyltransferases (DNMTs) re-establish 5mC patterns, maintaining methylation equilibrium [11] (Fig. 1A).

**Fig. 1.**
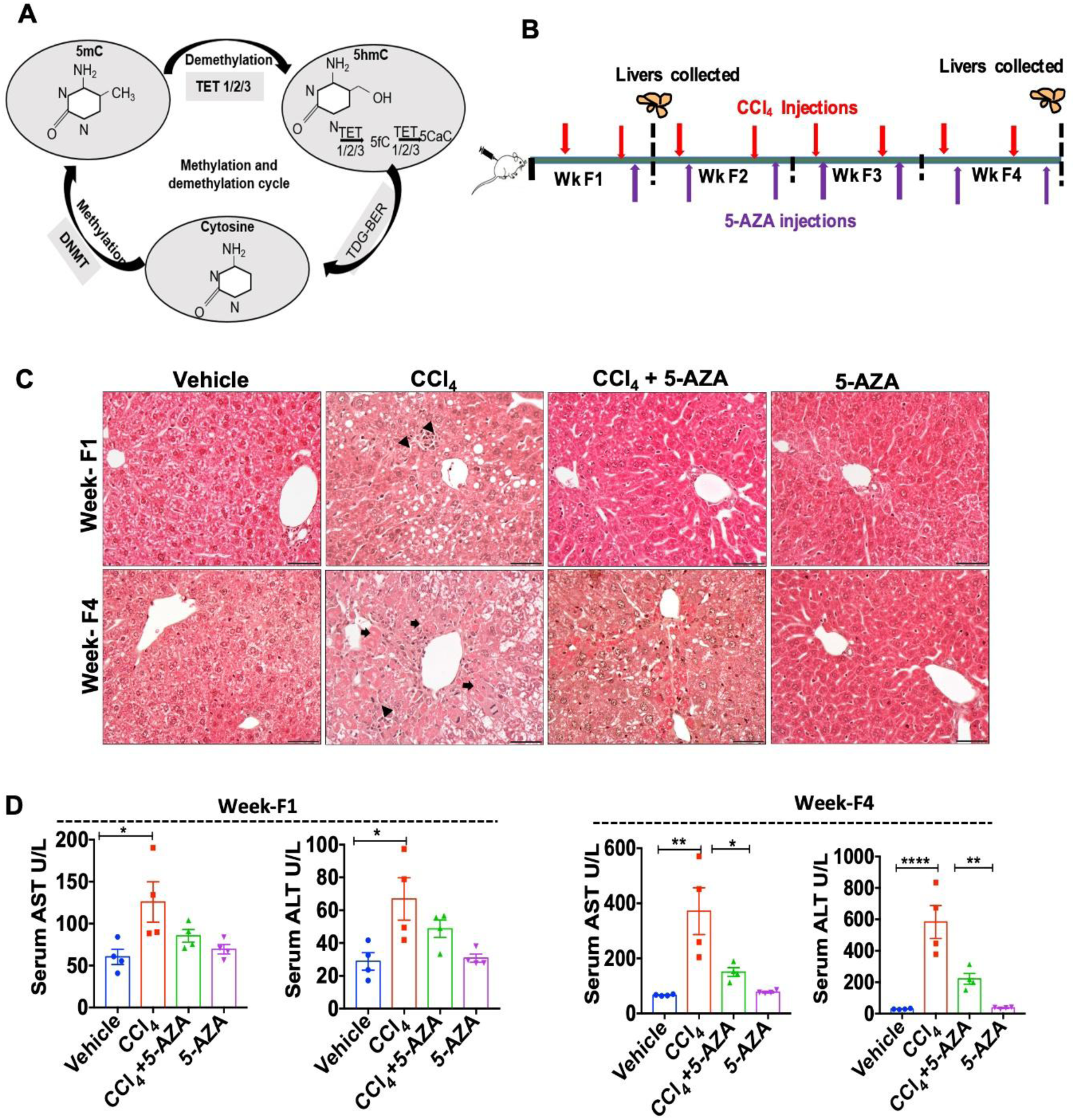
5-AZA treatment improved liver morphology and lowered serum AST and ALT activities during fibrosis phase. (**A**) DNA methylation-demethylation cycle: 5-methylcytosine (5mC) is oxidised by DNA demethylases TET1/2/3 to 5-hydroxymethylcytosine (5hmC) which is further oxidised to 5-formylcytosine (5fC) and 5-carboxylcytosine (5caC). The thymine DNA glycosylase (TDG) and the base excision repair (BER) pathway work together to replace these bases with unmethylated cytosine. This unmodified cytosine is methylated by DNA methyltransferase (DNMT) to regain the 5-methylcytosine (5mC). (**B**) Timeline for CCl_4_ and 5-AZA injections in a fibrosis progression model. (**C**) Representative images of H&E staining of liver sections from different groups; vehicle control, CCl_4_, CCl_4_+5-AZA and 5-AZA at fibrosis phase (Weeks F1& F4). Foci of cellular infiltrates (arrow heads), and hepatocytes with injured morphology (arrows) are shown. (**D**) Serum AST & ALT at Week F1 & F4 (n=4). Scale bars (50 µm). *P<0.05, ** P<0.01, **** P<0.0001 significance by one-way ANOVA with Turkey’s post test.

Aberrant regulation of this methylation-demethylation cycle has been increasingly linked to fibrosis with several studies highlighting alterations in DNMTs and TETs during liver fibrosis [6,12,13]. A global loss of 5hmC, together with variable expression of TET proteins, has been reported in different murine models of liver fibrosis [6]. Recent investigations have revealed that TET1 contributes to the progression of non-alcoholic fatty liver disease (NAFLD) and Alcohol liver disease (ALD) while TET3 acts as a determinant of fibrosis severity, underscoring the functional significance of DNA demethylation in hepatic pathology [14–16]. Epigenetic drugs that modulate DNA methylation offer a potential therapeutic avenue for reprogramming these aberrant changes. The DNMT inhibitors 5-Azacytidine (5-AZA, Vidaza) and 5-AZA-2′-deoxycytidine (5-AZAdC, Decitabine) are FDA-approved agents for hematological malignancies [17,18]. These are now being explored for repurposing in fibrotic diseases and preclinical studies have shown that 5-AZA and 5-AZAdC treatment attenuates fibrosis in the lung, heart and kidney [19–21]. Furthermore, 5-AZAdC was shown to block *in vitro* myofibroblast differentiation of rat liver HSCs [22]. However, these findings primarily attribute antifibrotic effects to DNMT inhibition, leaving the role of TET modulation largely unexplored.

An earlier study, demonstrated that 5-AZA either alone or in combination with Vitamin C not only inhibits DNMTs but also promotes active DNA demethylation by upregulating TET2 and TET3 in hepatocellular carcinoma (HCC) cells, leading to apoptosis [23, 24]. To our knowledge, targeting DNA demethylation as a therapeutic strategy in liver fibrosis has not been reported. In this study, we investigated whether pharmacological modulation of DNA hydroxymethylation could alter the course of fibrosis and recovery. Using both *in vivo* and *in vitro* models of liver fibrosis, we evaluated the effects of 5-AZA treatment on TET expression, global 5hmC levels, and fibrogenic responses. Our findings reveal that with 5-AZA treatment there is restoration of 5hmC and TET1 levels while TET3 levels are reduced with amelioration of fibrotic pathology. These results establish the functional relevance of TET-mediated hydroxymethylation in liver fibrosis and highlight this axis as a promising therapeutic target for epigenetic intervention.

## 2. Materials and methods

### 2.1 Animal studies

Mice from inbred colonies of C57BL/6 (male mice) maintained at animal facility of the Centre for Cellular and Molecular Biology (CCMB), Hyderabad, India was used after obtaining appropriate approvals from the Institutional Animal Ethics Committee. All experimental procedures were performed according to the approved guidelines for care and use of animals.

### 2.2 Mouse model for liver fibrosis and recovery

Male mice (8-9 weeks) were randomly segregated into four groups (n=4 each): Vehicle control, CCl_4_, CCl_4_+5-AZA and 5-AZA. Liver fibrosis was induced by intraperitoneal injections of CCl_4_ (0.6 mL/kg body weight) twice weekly for 4 weeks. Impact of 5-AZA was tested during fibrosis progression phase and recovery phase (Fig 1B,E).

In the fibrosis progressive model, mice in the CCl_4_+5-AZA group received 5-AZA (5 mg/kg body weight) twice a week, on the next day after CCl_4_ injection for 4 weeks (Week F1-F4). In the recovery model, mice in the CCl_4_+5-AZA group received 5-AZA after completion of 4 weeks of CCl_4_, and continued for another 4 weeks (Week R1-R4). 5-AZA control group received only 5-AZA and vehicle control group received only olive oil. Mice were sacrificed at Week F1 & F4 and R1 & R4 and liver tissue harvested. Blood was collected before the sacrifice and serum was used to measure Aspartate amino transferase (AST) Alanine amino transferase (ALT) activities using kits from Coral Clinical Systems, India, Pvt. Ltd.

### 2.3 Isolation and culture of Mouse Primary hepatic stellate cells

HSCs were isolated using the protocol of Maschmeyer et al [25]. In situ liver perfusion was performed sequentially with pronase E solution and collagenase P, digested and livers pooled from five mice were used to isolate the primary HSCs by 8% Nycodenz gradient (Fig. S1). Cells were cultured in Dulbecco’s Modified Eagles Medium (DMEM) with 10% FBS. On day 4, cells were treated with 5µM of 5-AZA for additional 72 h and harvested. Untreated cells at after initial 24h of culture and on Day 7 served as Day 1&7 controls respectively.

### 2.4 Cell Culture TGF-β treatment and transfection

LX-2 (human HSCs) were maintained in DMEM with 2% serum. At 70-80% confluence, cells were serum starved for 24h, treated with TGF-β (2.5 µg/ml) and 5-AZA (2.5 µM) either independently or in combination for another 48h and fixed for immunostaining or harvested for protein. Cells seeded in a 12-well plate were transfected with si-TET3 at 10pM (siRNA(h) and si-Scr (Control siRNA-A) (Santacruz biotechnologies, USA) using Lipofectamine RNAiMAX reagent (Invitrogen, USA) and protein isolated after 48h.

### 2.5 Histology Immunohistochemistry (IHC)and immunocytochemistry (ICC)

Paraffin embedded liver sections were stained for Hematoxylin and Eosin (HE) and Sirius Red (SR, Sigma-Aldrich, USA). IHC was performed using 5mC, 5hmC, α-SMA and TET1 antibodies (Abcam, USA). Visucyte HRP polymer (R&D biosystems, USA) was used as secondary followed by DAB staining. For fluorescent detection, anti-rabbit-alexa 594 (Molecular Probes, USA) was used as a secondary antibody. For ICC, 5hmC, α-SMA and COL-I primary antibodies (Abcam) and either anti-rabbit-Alexa 594/488 as secondary antibodies were used.

### 2.6 Imaging and measurements

Images were captured using Olympus inverted microscope (IX3-SSU, Tokyo, Japan) or Axioplan 2 imager (Zeiss, Germany). All the images within a comparison set were captured using identical exposure and gain settings. Total percent of 5mC and 5hmC positive nuclei were scored by IHC profiler software that classifies and counts individual nuclei as high/medium/low/negative based on the DAB colour deconvolution and reports the percentage of positive nuclei [26]. Image J was used to quantify the SR stained/α-SMA IHC sections and the fluorescence intensity of 5hmC in primary HSCs and LX-2 cells.

### 2.7 RNA isolation and real-time PCR

Total RNA from cultured primary stellate cells was isolated using the Trizol^TM^ reagent. Prime script first strand cDNA synthesis kit (Takara, Japan) was used for cDNA preparation. Roche Light cycler syber green master mix (Roche) was used for real time PCR and expression of *Hprt* was used to normalise gene expression. Primer sequence is given in Table S1.

### 2.8 Immunoblot assays

Total protein from the liver tissue and cultured cells was isolated with RIPA lysis buffer. Proteins were separated on SDS-gels, transferred on PVDF membranes and probed for TET1 (Abcam, USA), TET3 (Abcam/Novus, USA) α-SMA (Abcam, USA) COL I (Merck Millipore, USA) and GAPDH (Santacruz Biotechnologies, USA) antibodies. Detection was performed using ECL western blotting substrate.

### 2.9 Statistical analysis

All statistical analysis was done using GraphPad Prism (ver. 7.0). Data is represented as mean ± SEM. Significance was calculated by one-way ANOVA with Tukeys post test. P-values of *P<0.05/0.03,**P<0.002, ***P<0.0002, ****P<0.0001 were considered significant.

## 3. Results

### 3.1 5-AZA treatment improves liver morphology, lowers serum AST and ALT activities and reduces fibrosis

The efficacy of modulating DNA hydroxymethylation on mitigating fibrosis was tested in (a) Fibrosis phase-with ongoing injury and simultaneous 5-AZA injections (Fig. 1B) and (b) Recovery phase-After CCl_4_ withdrawal, 5-AZA injection is initiated (Fig. 2A). In the CCl_4_ group, by Week F1, H & E staining of the liver tissue revealed morphological features such as regions of damaged hepatocytes and presence of foci of infiltrated cells commonly seen in injured livers. While the livers of mice in the treated group of CCl_4_+5-AZA, at the same time point had lower foci of infiltrated cells and the tissue appeared less damaged compared to the CCl_4_ group (Fig. 1C). By Week F4, histological features consistent with hepatocyte injury were evident in livers of the injured group (CCl_4_), while, livers of mice in treated group (CCl_4_+5-AZA) at the same time point, did not show evidence of severe injury. Serum AST & ALT activities were significantly increased in the CCl_4_ group from weeks F1 to F4 as compared to vehicle control mice, while the serum AST & ALT activities were lower in the CCl_4_+5-AZA group at both weeks F1 and F4 (Fig. 1D).

**Fig. 2.**
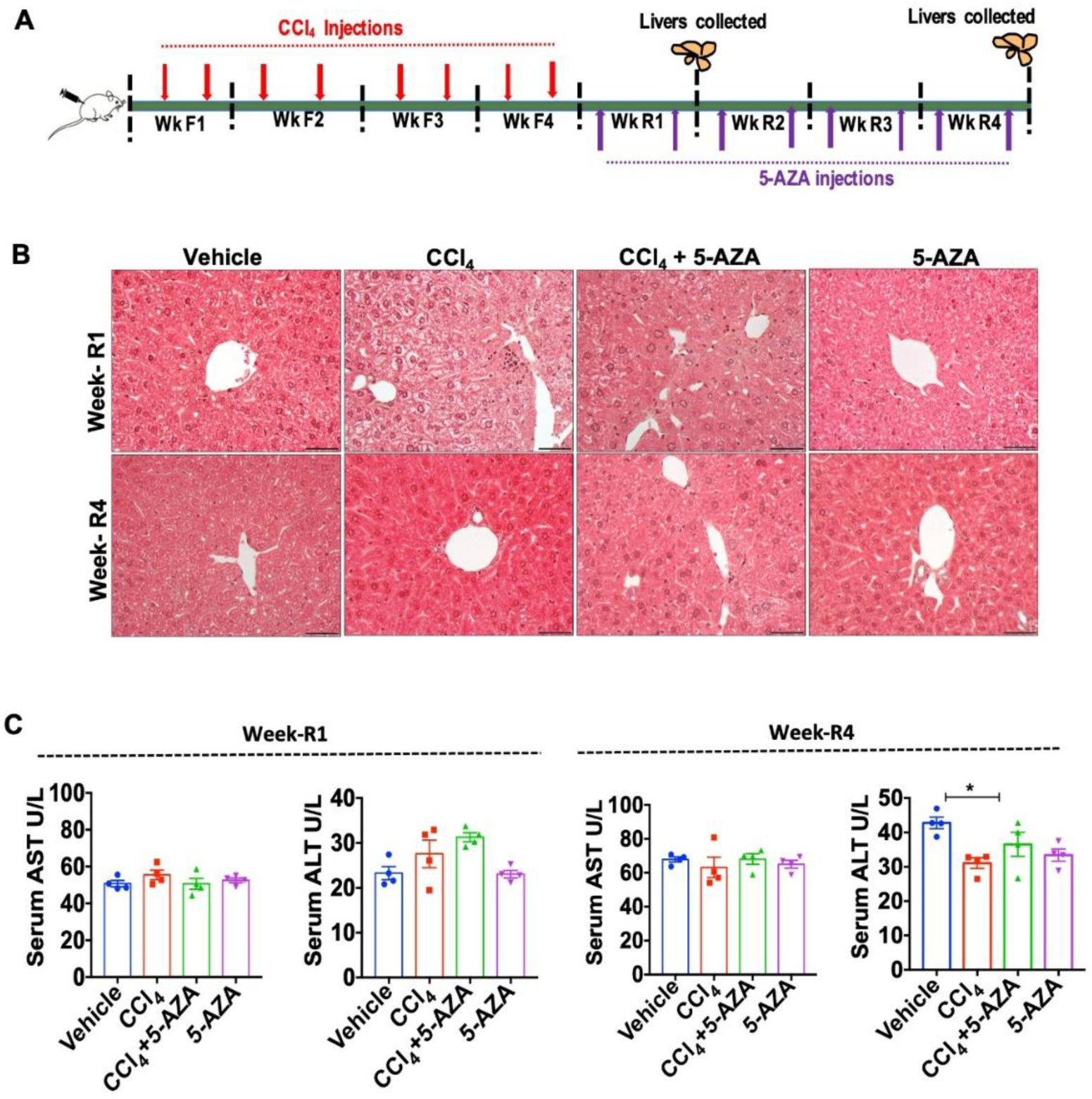
5-AZA treatment improves liver morphology during recovery. (**A**) Timeline for CCl_4_ and 5-AZA injections in a fibrosis progression and recovery model. (**B**) Representative images of H&E staining of liver sections from different groups; vehicle control, CCl_4_, CCl_4_+5-AZA and 5-AZA at recovery phase (Weeks R1 & R4). (**C**) Serum AST & ALT at Week R1 & R4 (n=4). Scale bars (50 µm). *P<0.05, ** P<0.01, **** P<0.0001 significance by one-way ANOVA with Turkey’s post test.

In the recovery phase, where 5-AZA treatment was initiated after CCl_4_ with drawl, a decrease in hepatocyte damage was observed in both untreated and treated livers by Week R1 but the foci of infiltrated cells were lower in the livers of the CCl_4_+5-AZA group (Fig. 2B). By Week R4, livers of both the CCl_4_ and CCl_4_+5-AZA groups recovered morphologically and enzyme levels also returned to normal (Fig 2C). No adverse changes in morphology or serology was seen in the mice livers that received 5-AZA alone during both fibrosis and recovery phases.

The extent of fibrosis, as measured by Sirius red staining (SR) and quantification, showed a progressive increase in collagen from Week F1 to Week F4, with bridging fibrosis, typical of CCl_4_ induced hepatotoxicity (Fig. 3A). In the livers of CCl_4_+5-AZA group, collagen was lower and there was no evidence of bridging fibrosis (Fig. 3A), suggesting collagen deposition is inhibited by 5-AZA treatment. During recovery, a decreasing trend in total collagen compared to the injury phase was clearly visible in the livers of both CCl_4_ and CCl_4_+5-AZA groups, however the decrease was significant in the CCl_4_+5-AZA group at Week R1 suggesting 5-AZA treatment enhanced the early recovery process (Fig. 3B).

**Fig. 3.**
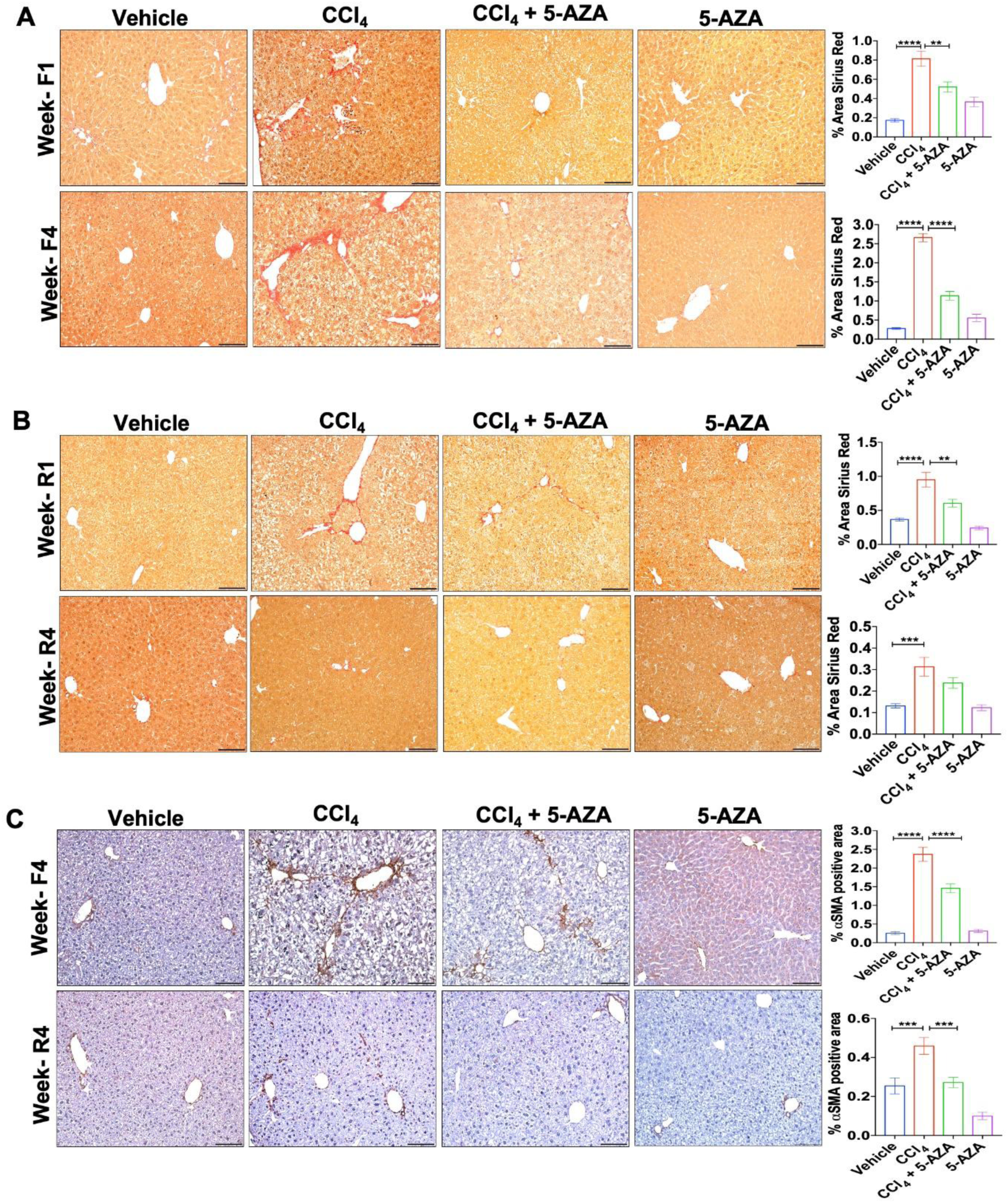
Fibrosis is reduced in CCl_4_ injured livers treated with 5-AZA. (**A, B**) Representative Sirius red stained images of liver tissue sections; (**A**) Week F1 & F4 during fibrosis and Sirius red quantification (**B**) Week R1 & R4 during recovery and SR quantification (graphs represent mean of 25-30 fields from n=4 mice livers/group) (**C**) IHC of liver tissues at Weeks F4 & R4 showing *α*-SMA staining (graphs represent mean of 25-30 fields from n=4 mice livers/group). Scale bars (100 µm). *P<0.03, ** P<0.002, *** P<0.0002, **** P<0.0001 significance by one-way ANOVA with Tukey’s posttest.

α-SMA; a marker for HSC activation when was assessed at Week F4 & Week R4 followed the same trend as collagen with significant decrease of α-SMA in the CCl_4_+5-AZA group as compared to the CCl_4_ group (Fig. 3C). At Week R4 as well, α-SMA was significantly lower in CCl_4_+5-AZA group when compared to the CCl_4_ group. Mice receiving 5-AZA alone did not show any significant changes in collagen or α-SMA.

### 3.2 An increase in TET associated hydroxymethylation (5hmC) is observed in fibrotic livers of mice treated with 5-AZA

Liver tissue sections stained for 5mC and 5hmC were quantified using the IHC profiler that counts individual nuclei as high/medium/low/negative positive based on DAB color deconvolution and reports the percentage of positive nuclei in each section. Immunohistochemical staining and quantification for 5mC (methylation mark) and 5hmC (hydroxymethylation mark) showed variations in the global methylation and hydroxymethylation status of liver at different stages of injury and recovery. The representative images of staining during fibrosis and recovery and the percentage of total positive nuclei scored by IHC profiler is shown (Fig 4 and 5). A detailed sub-grading of the percentage of high, medium and low positive nuclei and the microscopy images at lower magnification are shown in Figs. S2-S4.

**Fig. 4.**
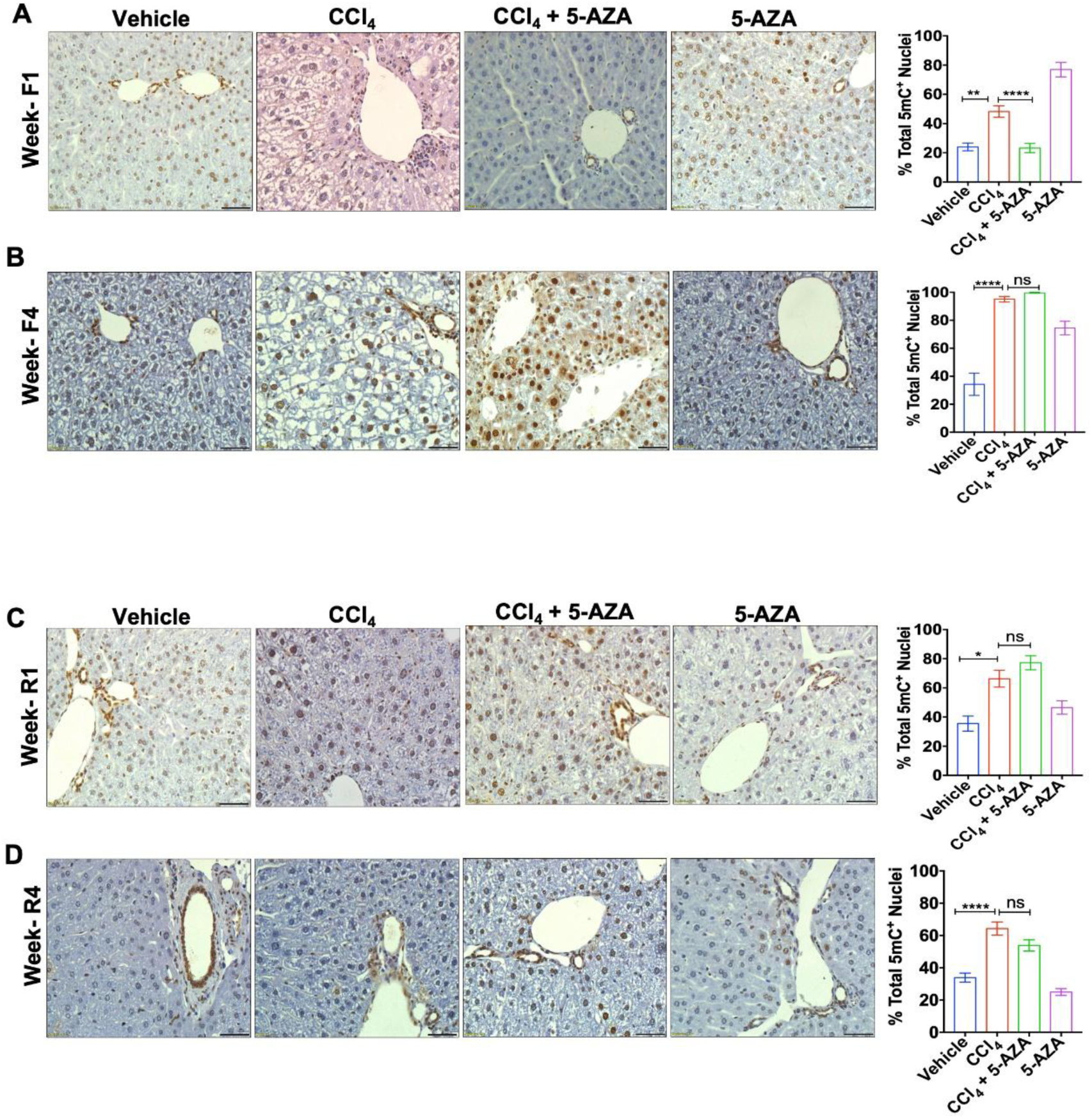
5-methyl Cytosine (5mC) levels are lower in CCl_4_ injured livers treated with 5-AZA during early phase of fibrosis. (**A, B**) Representative images of IHC staining of 5-methyl Cytosine (5mC) at week F1 & F4 and (**C, D**) Week R1 & R4. Methylation status was assessed by scoring the total percentage of 5mC-positive (brown) nuclei using the IHC profiler software (graphs represent mean of 25-30 fields from n=4 mice livers/group covering the entire section). Scale bars (50 µm). Error bars are mean with SEM. ns-non significant, * P<0.03, ** P<0.002, **** P<0.0001 significance by one-way ANOVA with Tukeys post test.

**Fig. 5.**
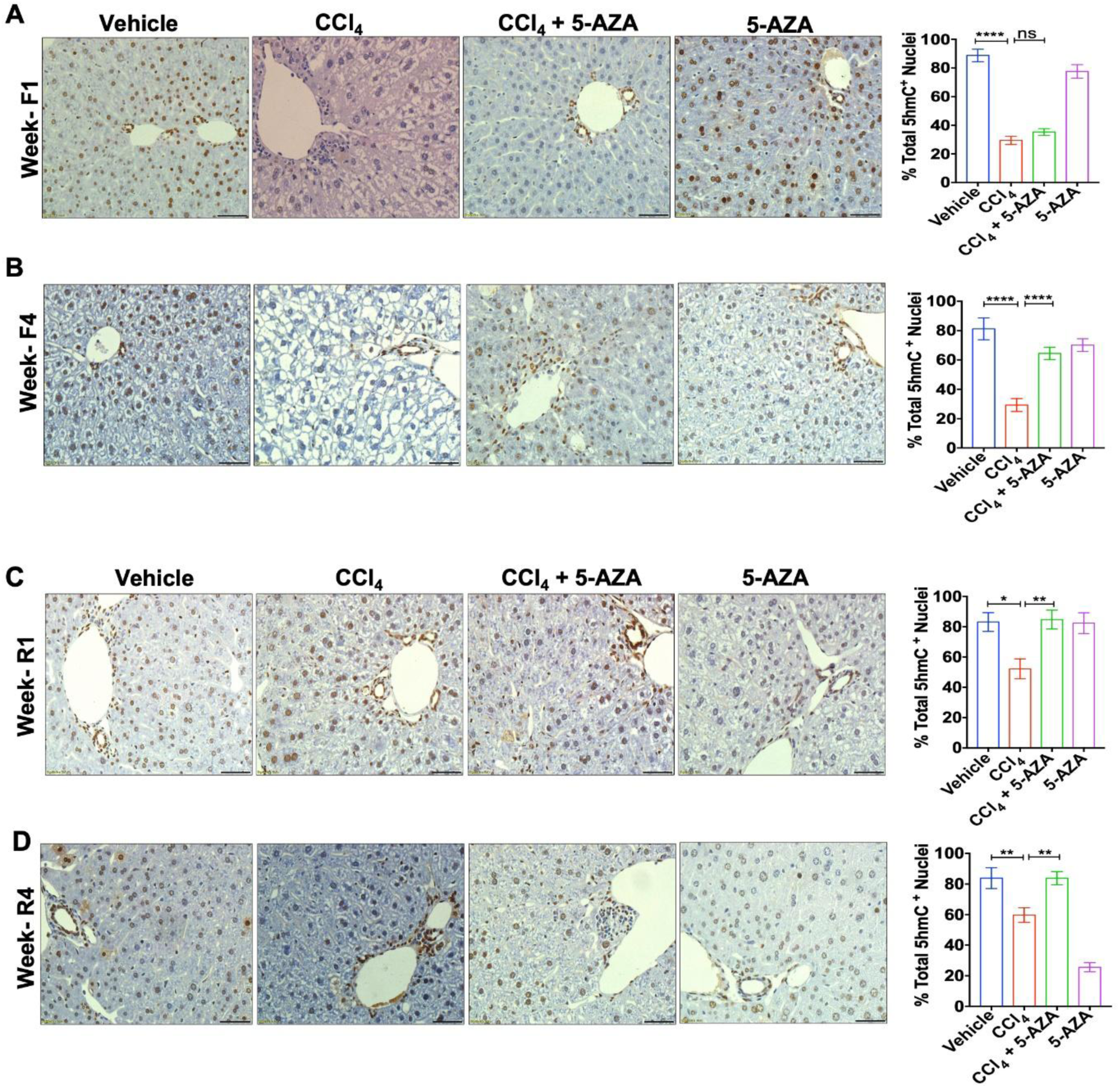
5-hydroxy methyl cytosine (5hmC) levels improved significantly in CCl_4_ injured livers treated with 5-AZA. (**A, B**) Representative images of IHC staining of 5-hydroxy methyl cytosine (5hmC) at Week F1 & F4 and (**C, D**) Week R1 & R4. Hydroxymethylation status was assessed by scoring the total percentage of 5hmC positive (brown) nuclei using the IHC profiler software (graphs represent mean of 25-30 fields from n=4 mice livers/group covering the entire section). Scale bars (50 µm). Error bars are mean with SEM. ns-non significant, * P<0.03, ** P<0.002, **** P<0.0001 significance by one-way ANOVA with Tukeys post test.

In the normal livers (vehicle control), 20-25% of the nuclei are 5mC positive (Fig. 4A). During fibrosis there is a progressive and significant increase in the % 5mC positive nuclei in the CCl_4_ injured livers from 50% at week F1 to 90% at week F4 (Fig. 4A & B), confirming increased methylation during fibrosis. In the livers of CCl_4_+5-AZA group, the % of 5mC positive nuclei were low at week F1, similar to the vehicle control, however by week F4, the numbers of 5mC positive nuclei in the livers of both CCl_4_ and CCl_4_+5-AZA group were similar, showing an increase in methylation (Fig. 4A &B). During recovery (weeks R1 and R4), in comparison to the livers of the vehicle control group, the % 5mC positive nuclei were higher in the livers of the CCl_4_ and CCl_4_+5-AZA group. However the % 5mC positive nuclei in both these groups was lower in the recovery phase as compared to Week F4 of fibrosis suggesting a spontaneous reduction 5mC levels during the recovery (Fig. 4 C & D). No significant difference in 5mC levels in the livers of 5-AZA alone group was noted at any time point.

On contrary, assessment of 5hmC status of the livers of vehicle control group shows, 90% 5hmC positive nuclei, indicating a highly hydroxymethylated status (Fig. 5). In the livers of the CCl_4_ group, these levels dropped drastically to 25% by week F1 and continued to stay low. In mice livers of CCl_4_+5-AZA group, the % 5hmC positive nuclei were low at week F1 but by week F4 there was restoration of restoration by increase in 5hmC nuclei to 60 % (Fig. 5 A & B). The livers of mice receiving 5-AZA alone did not show any significant changes in 5hmC status in this fibrosis phase (Fig. 5 & B).

During recovery phase, total % 5hmC positive nuclei in the CCl_4_ group was low at 45-50% and a minimal increase from week R1 to R4 was seen (Fig. 5 C & D). In the CCl_4_ group that received 5-AZA (CCl_4_+5-AZA group), 5hmC was restored to levels comparable to the livers of the vehicle control group (Fig. 5C & D), indicating the positive effect of 5-AZA treatment in restoring 5hmC status of the liver.

These results suggest a progressive increase in global methylation and a corresponding decrease in hydroxymethylation during fibrosis progression. 5-AZA treatment does not appear to substantially affect global 5mC levels, but shows a positive effect on improving the global 5hmC levels during both fibrosis progression and recovery.

### 3.3 5-Aza treatment restores TET1 expression and lowers TET3 expression

To assess contribution of TETs to the changes in hydroxymethylation, TET1 and TET3 were analysed at peak of fibrosis (Week F4) and recovery (Week R4) (Fig. 6A,B). A significant decrease in TET1 levels in livers of CCl_4_ group was observed compared to the vehicle control group. However, the TET1 levels were increased in the livers of CCl_4_ group that received 5-AZA (CCl_4_+5-AZA group) (Fig. 6A), furthermore, it remained unchanged in the group receiving 5-AZA alone. On the contrary, TET3 levels increased in livers of CCl_4_ group as compared to vehicle control group and a decrease was observed in CCl_4_+5-AZA group (Fig. 6A). During the recovery phase too, protein level of TET1 showed a similar trend as during the injury phase, however, levels of TET3, did not vary significantly (Fig. 6B).

**Fig. 6.**
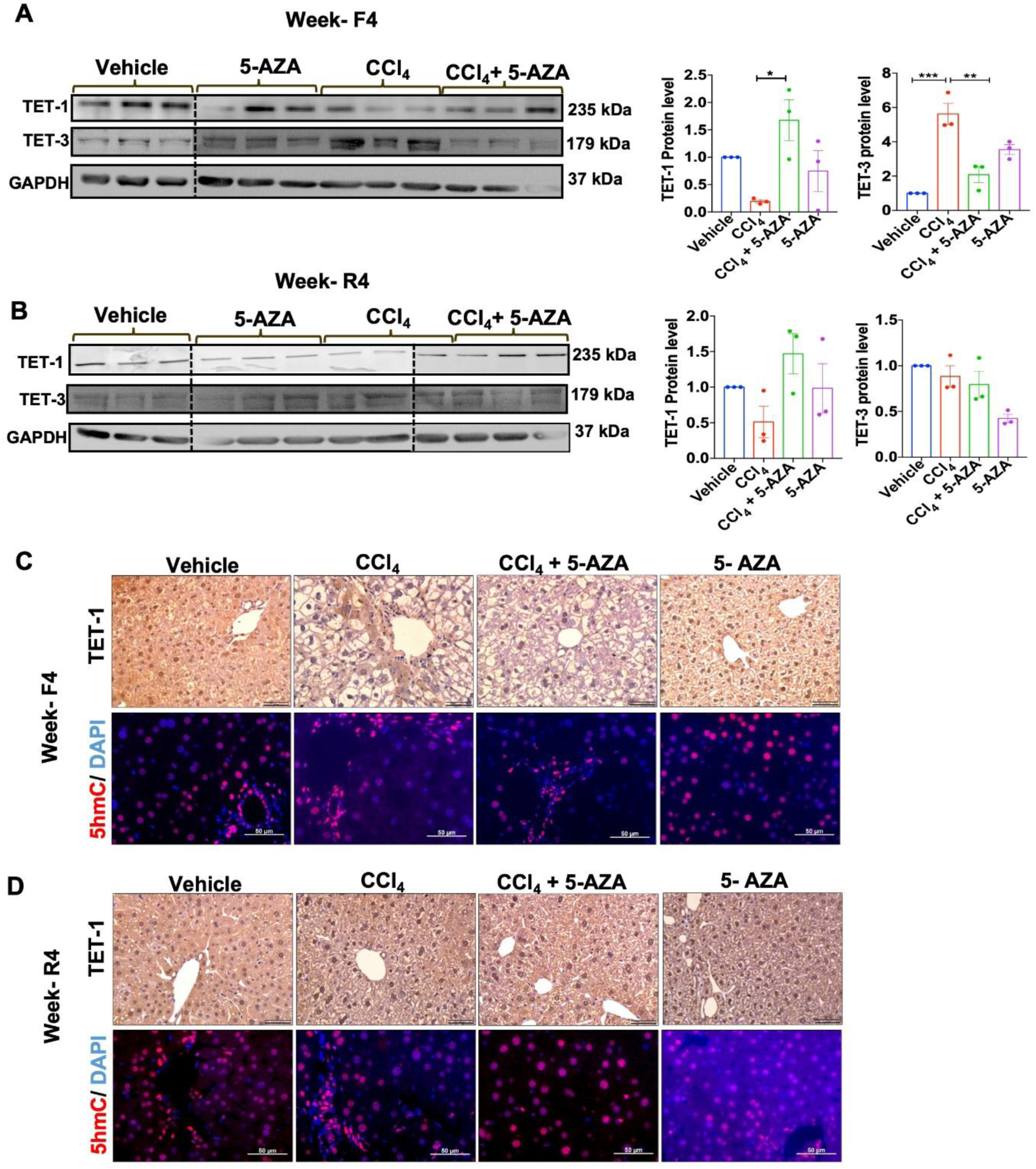
5-AZA treatment leads to enhanced TET1 and downregulated TET3 in fibrotic mouse livers. (**A**) TET1 & TET3 immunoblot from total liver protein at Week F4 (n=3). (**B**) Tet1 & Tet3 immunoblot from total liver protein at week R4 (n=3). (The dotted lines separates different blots grouped into the image). Protein levels are quantified using GAPDH as the loading control. **(C, D**) Representative images of IHC of TET1 (brown) & 5hmC (red-fluorescence) from different groups at Week F4 & R4. TET1 (25X magnification) & 5hmC (40X magnification). Scale bar-50 µm. Error bars are mean with SEM. *P<0.03, **P<0.002, *** P<0.0002, significance by one-way ANOVA with Tukey’s post test.

### 3.4 TET1 expression in the liver tissue corelates with 5hmC levels

Immunostaining for TET1 and 5hmC by individual staining, revealed that TET1 is predominantly observed in the hepatocytes (based on morphology of the cells and larger nucleus) (Fig 6 C) and its expression correlates with 5hmC levels (Fig 6 D). In CCl_4_ group at Week F4, this expression of TET1 and 5hmC (Fig 6 C&D) is reduced drastically as compared to the vehicle controls. While, in the 5-AZA treated fibrotic mice livers (CCl_4_+5-AZA group), an increase in both TET1 and 5hmC was observed.

During recovery (Week R4), there was no appreciable difference in TET1 and 5hmC levels in the CCl_4_ and CCl_4_+5-AZA groups (Fig. 6 E & F). Since, TET1 is predominantly observed in the nuclei consistent with hepatocyte morphology, and which form more than 50% of the liver cells, loss of 5hmC in these cells is reflected as a global and strong decline in 5hmC. Increased TET1 levels with 5-AZA treatment in the fibrotic livers, thus suggest a correlation between restored TET-1expression and increased global 5hmC status.

### 3.5 Reduced Tet3 expression by 5-AZA treatment in primary mouse HSCs

The prominent presence of 5hmC in fibrotic zones in the livers of the CCl_4_ group prompted us to further check for TETs expression in HSCs. HSCs freshly isolated from the livers of healthy mice and cultured (Fig 7A,B) were quiescent as identified with lipid droplets and autofluorescence (Fig. 7B). Once in culture, HSCs acquire fibroblastic morphology, lose their lipid droplets and autofluorescence gradually (Day 4 & Day 7) (Fig. 7B). Immunostaining for 5hmC, showed low levels in Day 1 control that increased by Day 7 in culture. Further upon 5-AZA treatment, the 5hmC levels were found to be lowered in these cells by Day 7 (Fig 7C). Upon assessing the changes in expression, an increase in *Tet1* & *Tet 3* (Fig 7D) was observed by Day 7. However, in the cells treated with 5-AZA, only *Tet3* expression was found to be reduced significantly (Fig. 7 D).

**Fig. 7.**
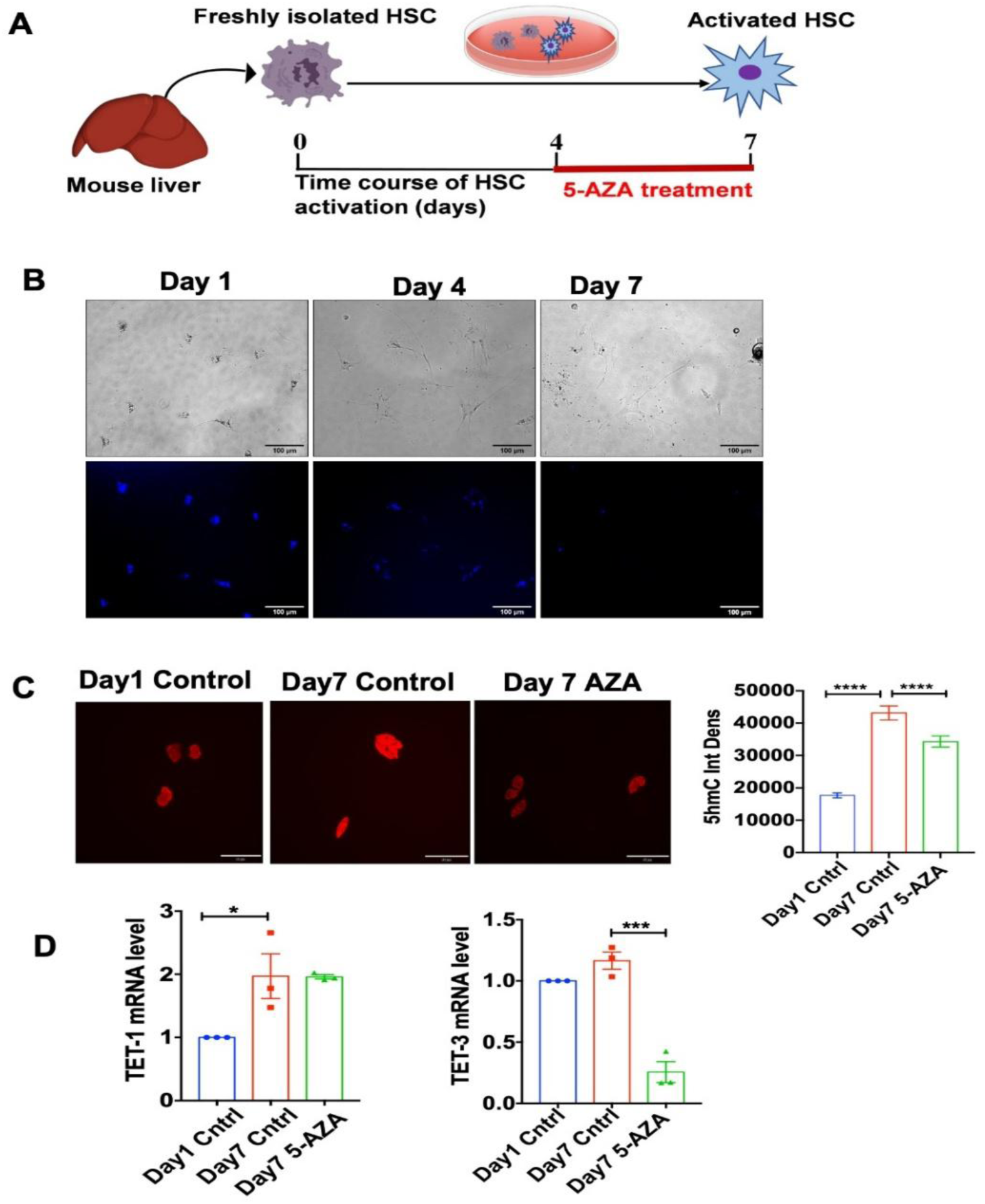
Decrease in 5hmC levels correlates with downregulated TET3 in 5-AZA treated primary HSCs. (**A**) Schematic of freshly isolated mouse primary stellate cells in culture and treatment with 5-AZA. (**B**) Primary HSCs in culture depicting changes in morphology of cells and Vitamin A fluorescence from day 1 to 7. (**C**) Cells in culture at day 4, treated with 5-AZA, fixed on day 7 and stained for 5hmC. Staining intensity quantified with Image J and shown as 5hmC Int Dens (mean of 140 nuclei for each group; n=3) (25X magnification; Scale bar-50 µm). (**D**) qPCR showing *Tet*1 & *Tet*3 levels in 5-AZA treated primary HSCs at day 7 compared to untreated day 1 and 7 controls (n=3). Error bars are mean with SEM. *P<0.03, P<*** P<0.0002, **** P<0.0001 significance by one-way ANOVA with Tukey’s post test.

### 3.6 LX-2 cells treated with 5-AZA show reduced TET3 correlating with reduced COL1 and α-SMA expression

To study the effect of 5-AZA on the expression of fibrosis markers in the stellate cells, we used the human hepatic stellate cell line LX-2. Cells treated with TGF-β for fibrosis induction show upregulated α-SMA and COL1 that were reduced in the presence of 5-AZA, while cells treated with 5-AZA alone showed no significant changes (Fig. 8A & B). Quantification of 5hmC, showed significantly higher 5hmC levels in TGF-β treated LX-2 cells, as compared to control (Fig. 8C). Upon treatment with 5-AZA a significant decrease in 5hmC was observed (Fig. 8C). Further, TET3 protein levels also correlated with these changes in 5hmC levels (Fig. 8D), suggesting 5-AZA treatment reduces TET3 and contributes to lower 5hmC levels with a corresponding decrease of fibrogenic markers in HSCs.

**Fig. 8.**
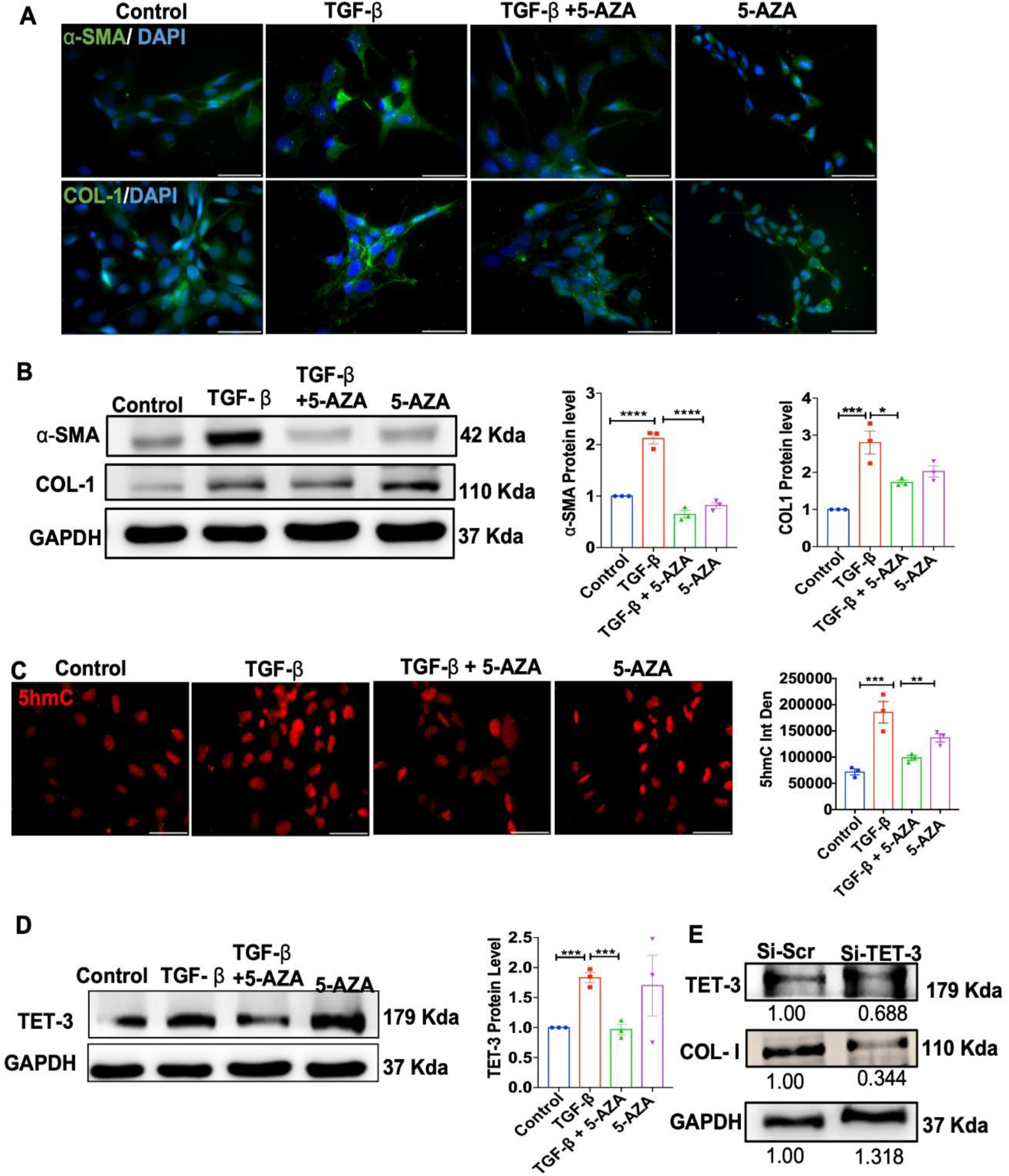
LX-2 cells treated with 5-AZA show reduced COL-I, α-SMA and 5hmC with a decrease in TET3. LX-2 cells from different treatment groups; untreated control, TGF-β, TGF-β+5-AZA and 5-AZA in culture for 48h were fixed for immunostaining and protein extracted for immunoblot. (**A**) Representative images of immunostaining for α-SMA & COL-I. (**B**) Immunoblot for α-SMA & COL-I (n=3). (**C**) Representative images of Immunostaining for 5hmC and quantification of staining intensity (n=3; mean intensity of total 300 nuclei/group quantified). (**D**) Immunoblot for TET3 and quantification (n=3). (**E**) knockdown of TET3 using si-TET3 in normal LX-2 cells shows lower TET3 expression and a corresponding decrease in COL-I. (Microscopy images are 25X magnification; scale bar-100 µm) Error bars are mean with SEM. **P<0.002, *** P<0.0002 **** P<0.0001 significance by one-way ANOVA with Tukey’s post test.

Further, LX-2 cells were transfected with si-TET3 to downregulate TET3 and immunoblotting confirmed a partial (∼31%) reduction in TET-3 levels relative to si-Scr control. A corresponding decrease in COL-I correlating with the decreased TET3 in the si-TET3 transfected LX-2 cells suggests a potential TET3 mediated regulation of COL-I (Fig. 8E).

## 4. Discussion

Liver fibrosis represents a maladaptive wound-healing response that underlies nearly all forms of chronic liver injury. Identifying strategies that can halt or reverse fibrosis at an early stage remains a major therapeutic challenge. Although numerous pharmacological candidates are under clinical investigation, effective antifibrotic therapies are still lacking. Increasing evidence suggests that targeting global regulatory mechanisms, such as epigenetic remodelling, could provide a unifying and durable therapeutic approach [27].

Epigenetic modulation of fibrogenesis was first demonstrated by Zeybel et al., who showed that inhibition of histone methyltransferase by 3-deazaneplanocin markedly attenuated liver fibrosis [28]. Among DNA-targeted epigenetic modulators, the DNA methyltransferase (DNMT) inhibitors 5-AZA and 5-AZA-2′-deoxycytidine (5-AZAdC) have been reported to alleviate fibrosis in several organs, including the lung, heart and kidney, through DNMT inhibition [19–21]. The reduction in α-SMA and COL-I expression observed in our study substantiates attenuation of HSC activation, corroborating antifibrotic outcomes seen in these earlier reported studies.

Sajadian et al. first demonstrated that in hepatocellular carcinoma cells, 5-AZA not only inhibited DNMT activity but also enhanced hydroxymethylation through upregulation of TET enzymes [23,24]. Our findings extend this paradigm to fibrotic liver disease by showing that 5-AZA treatment is accompanied by altered TET expression and global 5hmC levels, suggesting coordinated changes in the DNA methylation and hydroxymethylation pathways resulting in amelioration of the fibrotic pathology.

Previous studies have shown that promoter-specific hypomethylation of profibrotic genes and hypermethylation of genes involved in extracellular matrix degradation occur during HSC activation [4]. Pharmacological DNMT inhibition is shown to reduce HSC activation; for example, activated HSCs treated with 5-AZAdC exhibit lower α-SMA and COL-I levels, linked to DNMT1 inhibition [29]. In line with these reports, we observed reduced DNMT1 expression in fibrotic livers treated with 5-AZA (Fig. S5), suggesting that modulation of both methylation and hydroxymethylation contributes to the antifibrotic phenotype.

A major strength of this study lies in documenting the hydroxymethylation dynamics at two defined time points during both fibrosis and recovery. Previous studies primarily showed extensive epigenetic remodelling with HSC activation, accompanied by 5hmC loss that was assessed at a single time point after fibrosis was established using total genomic DNA [6]. In our study, by contrast, using immunostaining of the liver tissue, combined with *in vitro*confirmations in freshly isolated murine HSCs, and human HSCs (LX-2 cells), we demonstrate stage associated 5hmC changes and provide complementary evidence for hydroxymethylation alterations during fibrogenesis in the two major hepatic cell types. The decrease in 5hmC observed in the total liver tissue during fibrosis was restored with 5-AZA treatment; conversely, the increase in 5hmC in the isolated HSCs declines with 5-AZA treatment, which might reflect a cell-autonomous response specific to the isolated culture system.

The loss of 5hmC is paralleled by reduced TET1 in the whole liver (immunoblot and IHC) where hepatocytes constitute the dominant cell type. TET1 is implicated in regulating lipid metabolism and inflammatory signalling in liver disease models such as NAFLD, ALD and HCC [14,15,30]. Recent evidence confirms TET1 expression in the hepatocytes, which is downregulated in steatosis-induced HepG2 cells. Reduced TET1 in the human hepatocyte-like cell line OUMS29 has also been shown to promote apoptosis [14,15]. In our study, the restoration of TET1 protein levels by 5-AZA treatment correlated with the recovery of 5hmC levels, prominently observed in nuclei consistent with hepatocyte morphology (co-staining with a hepatocyte-specific marker to quantitatively confirm this cell-type attribution was not performed). While these results showed an association between 5-AZA treatment and restoration of TET1, loss- or gain-of-function studies of TET1 in the hepatocytes are required to establish whether 5-AZA’s effect on 5hmC is causally mediated via TET1 restoration. A recent study shows that TET1 also directly regulates HSC activity and inhibits fibrogenesis [31]. Together, these findings point to a broader role for TET1 in liver epigenetic regulation, beyond its effect on 5hmC alone.

TET3 has been documented as a fibrosis-promoting factor by Xu et al., [16]. In agreement with these findings, our study shows elevated TET3 expression during CCl₄-induced fibrosis that was reversed in mouse livers treated with 5-AZA, suggesting a regulatory cross-talk between methylation and hydroxymethylation pathways. Functional analyses in mouse and human HSCs demonstrated that TET3 downregulation was accompanied by reduced 5hmC and suppression of profibrotic markers. Knockdown of TET3 by siRNA correlated with decreased COL-I; thus, assessing 5hmC levels in TET3 knock down cells in future studies could further substantiate a direct role for TET3 in regulating 5hmC in these cells. Nevertheless, our observations so far are consistent with previous reports identifying TET3 as a determinant of hepatic fibrogenesis and a potential biomarker for chronic liver disease [16, 33, 34].

Few studies have reported variable or even decreased TET3 expression in fibrotic livers [6,33]. Page et al., reported decreased TET3 expression during *in vitro* HSC activation and *in vivo* across different rat models of liver fibrosis, accompanied by reduction in global 5hmC. However, they also report no change in TET3 in human alcohol-related liver disease [6]. In the study by Liu et al., TET3 expression was reduced in fibrotic liver and overexpression of TET3 led to increased 5hmC and promoted apoptosis of activated HSCs [33]. In contrast, in the study by Xu et al., TET3 increased in both human and mouse fibrotic livers [16]. Our earlier work with hepatocellular carcinoma cells *in vitro* also demonstrated increased TET2/TET3 levels when treated with 5-AZA [24]. The apparent differences between these studies suggest that TET3 could exert locus-dependent, opposing effects on profibrotic versus antifibrotic genes in different fibrosis models and the stage of fibrosis such that the net phenotypic outcome depends on the target loci that dominate the given model.

An interesting observation in this study was an increase in TET2 transcript levels in the CCl_4_-injured fibrotic livers, which was reduced following 5-AZA treatment at week F4. In contrast, no change in expression was observed at week R4 (Fig. S6). These findings differ from those of Page et al. and Wang et al., who observed a decrease in TET2 in different fibrosis and NAFLD models [6,14], and from another study that found no change in TET2 levels in fibrotic livers [16]. TET2 is predominantly characterized for its role in hematopoietic development and malignancy where it is highly expressed in myeloid and lymphoid lineage cells; a more limited body of evidence also implicates TET2 in hepatic gluconeogenesis [35–37]. Since the transcript levels of TET2 was measured in the whole-liver tissue rather than in isolated hepatocytes or HSCs, this predominantly hematopoietic expression pattern raises the possibility that the increase observed at Week F4 does not solely reflect changes within resident liver cells, but could in part be driven by the greater proportion of infiltrating immune cells during active fibrosis [38]. Consistent with this interpretation, the absence of change in TET2 transcripts at Week R4 could correspond with the reduced immune cell infiltration that accompanies recovery [38]. TET2 expression in primary stellate cells did not show significant changes (Fig S7), arguing against a stellate-cell origin for the Week F4 increase, although a contribution of TET2 to the global hepatic 5hmC cannot be ruled out. Further validation via cell-type specific analysis of infiltrating immune cells from resident hepatocytes and HSCs, is therefore needed to resolve this.

The contrasting effects observed across these different fibrosis models could reflect differences in fibrosis stage, species (rat vs mouse), and the specific cell population examined. These variables may be decisive, pointing to context dependent roles for TETs that are possibly shaped by differential chromatin accessibility or cofactor availability.

The dose of 5-AZA used in this study is consistent with prior reports that used the same dosage (5mg/kg body weight) for epigenetic intervention in experimental cardiac remodelling, demonstrating antifibrotic effects with reduced 5mC [20]. In our study, the administered dose was well tolerated, with no overt clinical or histological abnormalities observed in any of the mouse groups that received 5-AZA. Treatment with 5-AZA in the fibrotic livers was further associated with improved serum AST/ALT activities, restoration of 5hmC and alterations in TET enzyme expression.

There are a few shortcomings with our study. We report only one dosage of 5-AZA; for further studies aimed at repurposing 5-AZA as an antifibrotic therapy, it will be essential to standardise dosage through detailed toxicity assessments. Most importantly, these data do not distinguish whether altered TET expression reflects a direct pharmacological effect of 5-AZA, an indirect consequence of DNMT inhibition, or a secondary effect of reduced fibrosis and altered cellular composition. To answer these questions with certainty, future studies with closer time points, using primary hepatocytes, HSCs and immune cells isolated from injured/fibrotic livers would allow us to delineate the cell types and specific genomic loci that drive the hydroxymethylation changes at TET promoters. Complementing this with TET1/TET3 knockdown or knockout studies, combined with locus-specific methylation/hydroxymethylation changes at the profibrotic and protective gene promoters and their downstream targets via bisulphite sequencing, would help link the epigenetic shifts we observed to specific functional genes. This represents an important direction for future investigation.

## 5. Conclusion

Our findings suggest that treatment of fibrotic livers with 5-AZA is associated with modulation of the hepatic DNA methylation–hydroxymethylation equilibrium through coordinated changes in TET1 and TET3 expression. This is accompanied by restoration of 5hmC homeostasis, and mitigation of fibrogenesis. Beyond their catalytic function, TET proteins may also influence chromatin structure and transcriptional regulation, suggesting additional, contributions to their antifibrotic effects. While repurposing of 5-AZA for liver fibrosis warrants further preclinical validation, our study provides evidence that pharmacological modulation of the TET-mediated hydroxymethylation axis represents a promising avenue for antifibrotic therapy.

## CRediT authorship contribution statement

**Venkatapathi Challa**: Methodology, data acquisition and processing. **Rachiraju Hema Sindhuja**: Methodology, data acquisition and processing. **Truptimayee Barik**: Methodology, data acquisition and processing. **Arushi Mansingh**: Methodology, data acquisition and processing. **Sivapriya Pavuluri**: Data analysis, Writing & review editing. **Thomas Michael Shiju**: Methodology, Data analysis; **Kalyani Pathuri**: Data analysis and review. **Chaturvedula Tripura**: Funding acquisition, Conceptualization, Original draft, Supervision, Resources, Project administration, Methodology, Writing – review & editing, Writing – original draft.

## Ethical approval

This study was performed in line with the principles of the declaration of Helsinki. The study was approved by the Institutional Animal ethics committee of the CSIR-CCMB, India (No. IAEC73/2019).

## Availability of Data and Materials

The supporting data is provided in the supporting information.

All the other data generated and analysed during this study are available from the corresponding author upon request.

## Acknowledgements

CT is thankful to Dr. Scot Friedman, Mount Sinai Medical School for sharing the LX-2 cells. CT gratefully acknowledges the support of Prof. Andreas Nussler in initiating this study.

## Supporting Information

**Table S1:** List of Primers used in the Study.

**Fig. S1:** Mouse primary HSCs isolation.

**Fig. S2:** Detailed scoring of 5-methyl cytosine (5mC) and 5-hydroxymethyl cytosine (5hmC) levels in mouse liver tissue at week F1 & F4 and Week R1 & R4

**Fig S3:** Representative lower magnification images of 5mC immunostaining.

**Fig S4:** Representative lower magnification images of 5hmC immunostaining.

**Fig. S5:** *Dnmt*1 expression in livers at Week F4 and R4.

**Fig S6**: *Tet2* expression in the livers at Week F4 and R4.

**Fig S7:** *Tet*2 expression in mouse primary stellate cells.

## ABBREVIATIONS

TET: Ten-eleven translocases
5-Aza: 5-Azacytidine
CCl_4_: Carbon tetra chloride
CLD: Chronic liver disease
HSC: hepatic stellate cells
5mC: 5methyl cytosine
5hmC: 5-hydroxymehyl cytosine
DNMT: DNA methyl transferase
TGF-β: transforming growth factor beta
5-AzadC: 5-aza-2-deoxycytidine
AST: Aspartate transaminase
ALT: Alanine transaminase

