## Supplementary files for "5-Azacytidine alleviates Liver Fibrosis via modulation of TET1 and TET3 levels and restoration of global 5hmC"

#### I. List of Primers used in the Study

| Gene | Seq text (5’-3’) |
|---|---|
| *Hprt*1 Forward<br>*Hprt*1 Reverse | ATCAGTCAACGGGGGACATA<br>CTAACCATTTTGGGGTGT |
| *Tet*1 Forward<br>*Tet*1 Reverse | CTCAGGTGGGTTTGCAGAAA<br>AGAACGGCTGTGGGAAGAG |
| *Tet*2 Forward<br>*Tet*2 Reverse | TGGCAGTACAGTGGTGGTCA<br>TGGGCAGCTCTCCTATCCTT |
| *Tet*3 Forward<br>*Tet*3 Reverse | AGTGGGTGATCCGAAGACAC<br>ATCATTGAGGCCACATCTCC |
| *Dnmt*1 Forward<br>*Dnmt*1 Reverse | AAGAATGGTGTTGTCTACCGAC<br>CATCCAGGTTGCTCCCCTTG |

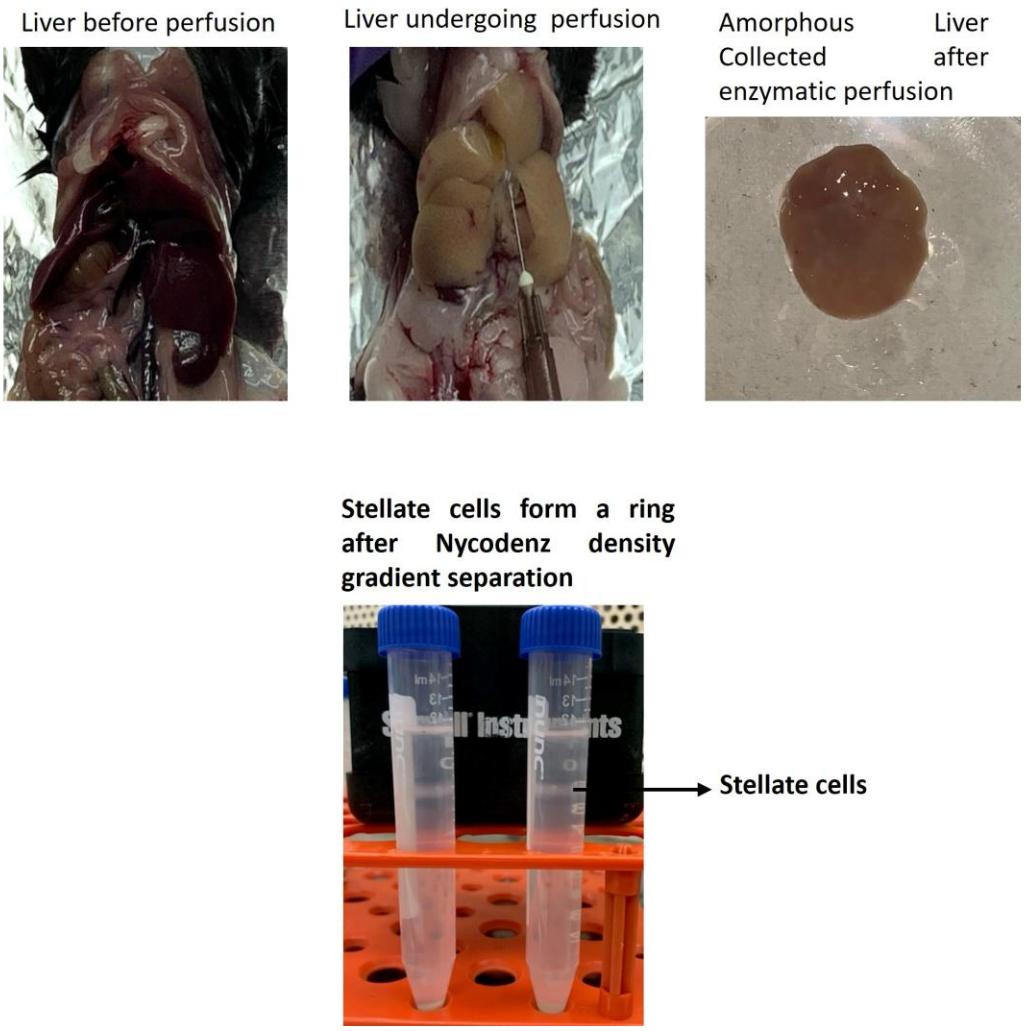


**Fig. S1: Mouse primary HSCs isolation**

Images of liver before and after continuous enzymatic perfusion depicting the changes in the liver appearance during the isolation of hepatic stellate cells.

The layer of stellate cells in the tube after Nycodenz density gradient separation is also shown

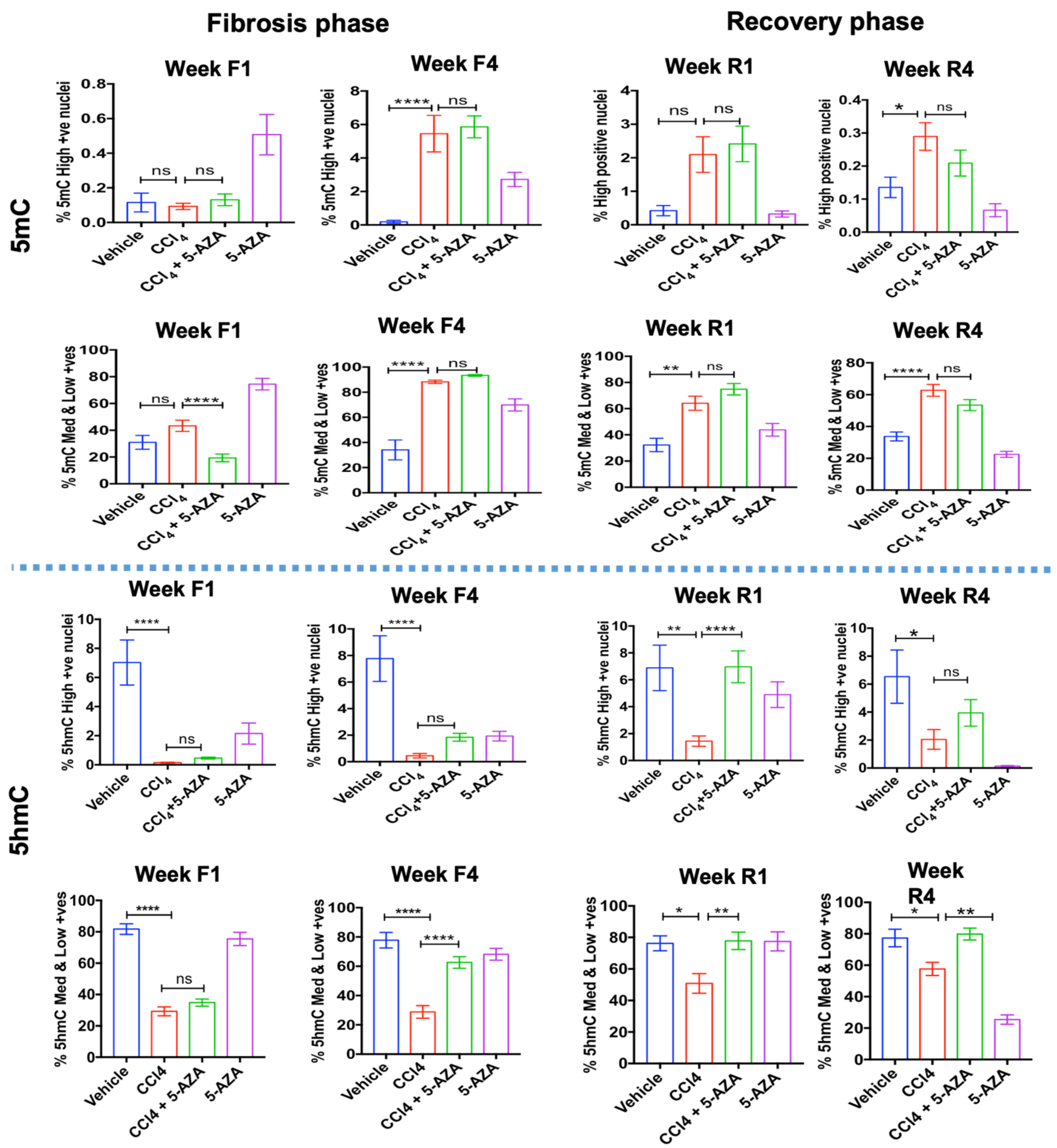


**Fig. S2: Detailed scoring of 5-methyl cytosine (5mC) and 5-hydroxymethyl cytosine (5hmC) levels in mouse liver tissue at week F1 & F4 and Week R1 & R4.** Methylation and hydroxymethylation status was assessed by scoring 5mC and 5hmC positive (brown) nuclei in the liver sections at Weeks F1 & F4 and Weeks R1 & R4. Quantification was by IHC profiler software as high, medium & low based on the DAB colour deconvolution (graphs represent mean of 25-30 fields from n=4 mice livers covering the entire section). Error bars are mean with SEM. ns-non significant, * P<0.03, ** P<0.002, **** P<0.0001 significance by one-way ANOVA with Tukeys post test.

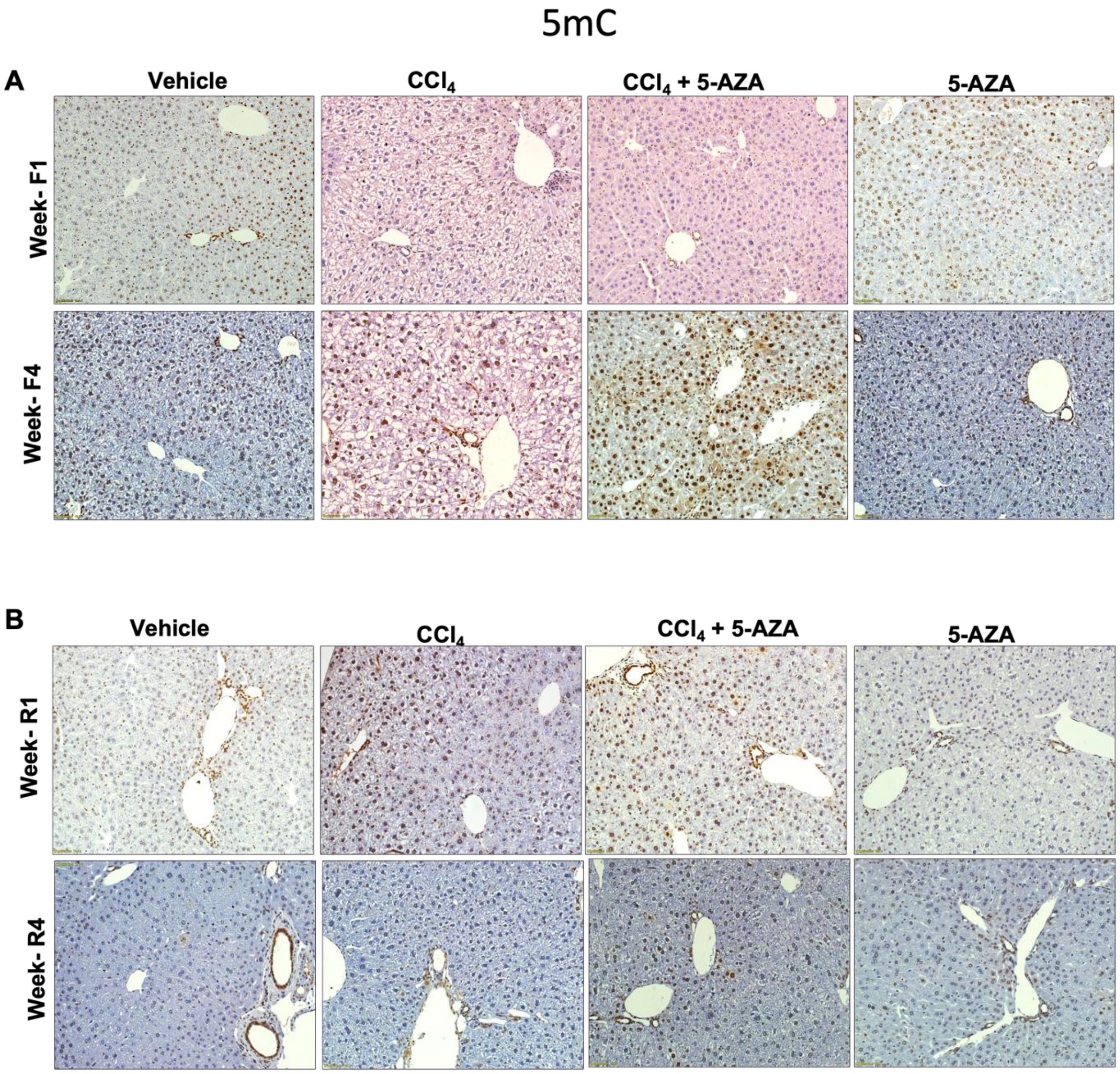


**Fig. S3: Representative lower magnification images of 5mC immunostaining.**

5 methyl cytosine (5mC) immunostaining at Week F1& F4 and Week R1 & R4 in the different groups (Vehicle control., $CCl_4$., $CCl_4$+5-AZA., 5-AZA). (20X objective; Olympus inverted microscope (IX3-SSU, Tokyo, Japan)

### 5hmC

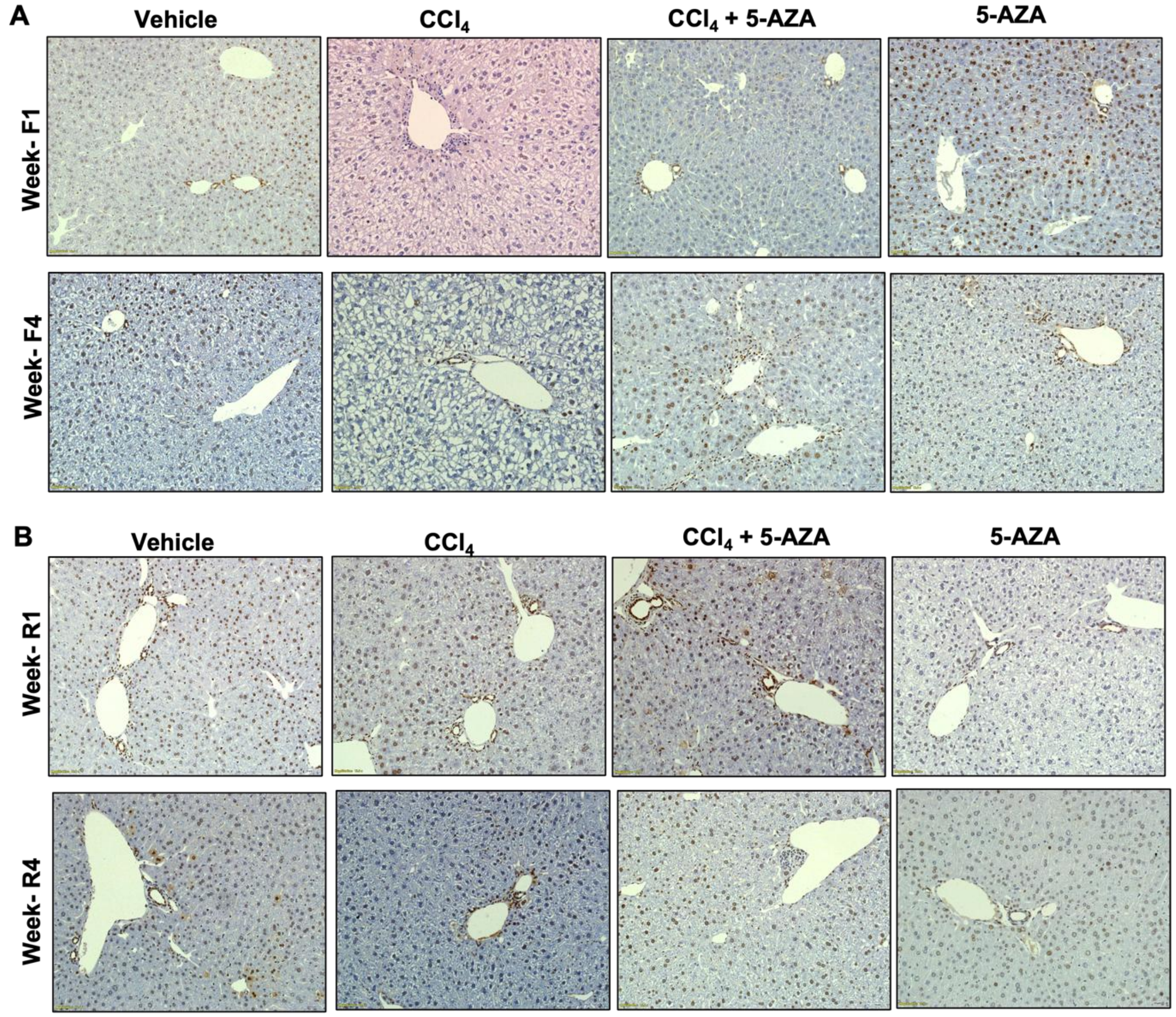


**Fig. S4: Representative lower magnification images of 5hmC immunostaining.**

5 hydroxymethyl cytosine (5hmC) immunostaining at Week F1& F4 and Week R1 & R4 in the different groups (Vehicle control., $CCl_4$., $CCl_4$+5-AZA., 5-AZA). (20X objective; Olympus inverted microscope (IX3-SSU, Tokyo, Japan)

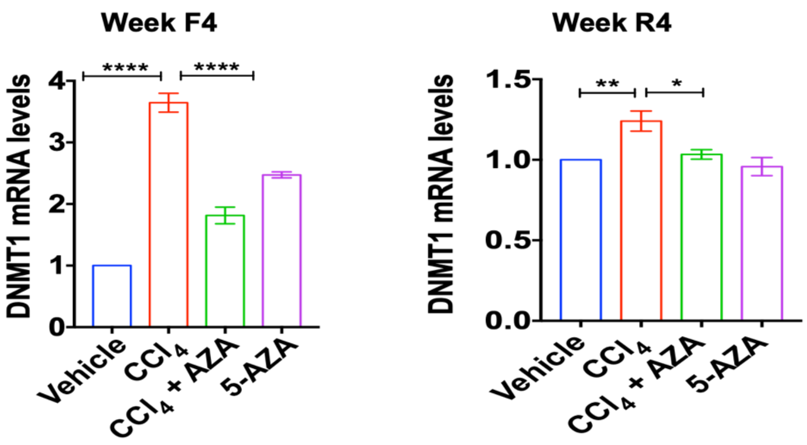


**Fig. S5: *Dnmt*1 expression in the livers at Week F4 and R4**

*Dnmt*1 expression was quantified by qPCR using *Hprt*1 as the house keeping gene in the different groups; Vehicle, $CCl_4$, $CCl_4$+5-AZA and 5-AZA (n=3). Error bars are mean with SEM. *P<0.03, P<** P<0.002, **** P<0.0001 significance by one-way ANOVA with Tukey's post test.

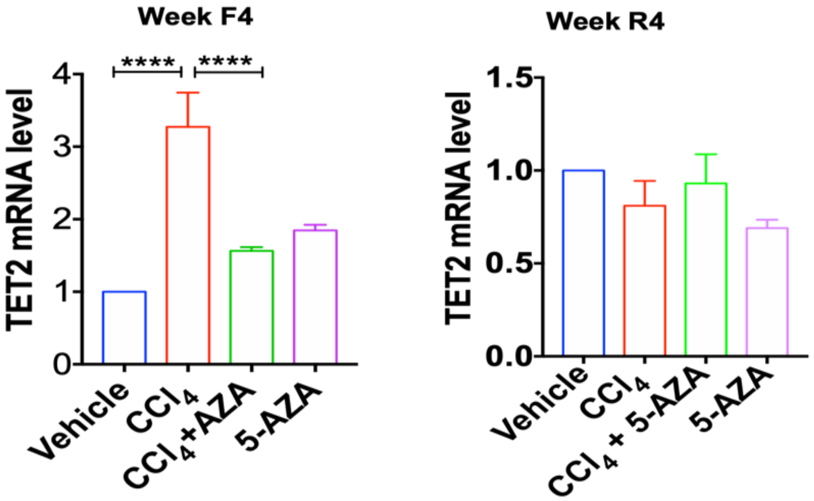


**Fig. S6: *Tet2* expression in the livers at Week F4 and R4**

*Tet2* expression was quantified by qPCR using *Hprt*1 at Week F4 and R4 by qPCR using *Hprt*1 as the house keeping gene in the different groups; Vehicle, $CCl_4$, $CCl_4$+5-AZA and 5-AZA (n=3). Error bars are mean with SEM. **** P<0.0001 significance by one-way ANOVA with Tukey's post test.

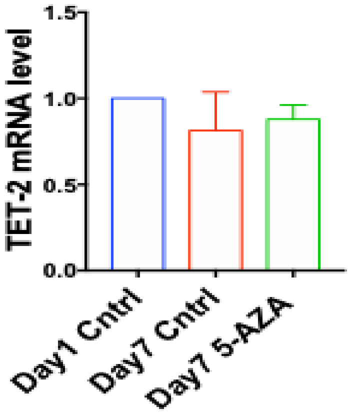


**Fig. S7: *Tet*2 expression in mouse primary stellate cells.**
Tet2 expression in the mouse primary stellate cells was quantified by qPCR with *Hprt*1 as the house keeping gene in 5-AZA treated primary HSCs at day 7 compared to untreated day 1 and 7 controls (n=3).
